# Plant domestication does not reduce diversity in root microbiomes

**DOI:** 10.1101/2025.01.13.632861

**Authors:** Alejandra Hernández-Terán, Ana E. Escalante, Maria Rebolleda-Gómez

**Author notes:** **Corresponding authors:** Alejandra Hernández-Terán, María Rebolleda-Gómez.

## Abstract

- Domestication has profoundly shaped the genetic makeup of numerous plant and animal species. While the effects of plant domestication at the genetic and phenotypic levels are well-documented, its impact on plant microbiome remains less understood.
- Two primary hypotheses have been proposed: 1) the reduction in microbial diversity resulting from the domestication process, and 2) the diminished ability of host plants to control their microbiomes.
- We conducted a meta-analysis of multiple crops, comparing the root microbiomes of domesticated plants and their wild relatives. Our results indicate that the effects of domestication are species-specific and context-dependent, with most domesticated plants exhibiting increased microbial diversity and more structured communities.
- Overall, this study provides evidence that plant domestication does not lead to a uniform reduction in microbial diversity or a consistently diminished ability of plants to influence their microbiomes.
- Based on these findings, we discuss new perspectives and the need for future studies incorporating native soils and host genetic variation in such experiments, analyzing diversity and microbiome function, and considering how root morphology might affect microbiome recruitment.

## INTRODUCTION

Domestication, the artificial selection of phenotypic traits useful to humans, has profoundly shaped human civilization and the genetic makeup of numerous plant and animal species (Purugganan & Fuller, 2009). The consequences of domestication at the genetic and phenotypic level are well-documented (Meyer *et al*., 2012). Typically, domestication events involve propagating only selected plant seeds, leading to genetic bottlenecks and, thus, a reduction in genetic diversity across the genome (Doebley *et al*., 2006). Phenotypically, convergent changes in certain traits, known as ‘domestication syndrome’, have been observed across diverse domesticated species (Alam & Purugganan, 2024). In plants, this syndrome is characterized by enlarged fruit sizes, increased apical dominance, and the loss of seed dispersal mechanisms and dormancy, among other traits (Doebley *et al*., 2006). Many of these traits are the target of selection, but many others are not. Those non-target traits can evolve through correlations (genetic linkages or pleiotropy) with the target traits (e.g. reduction of seed shattering, uniform flowering, and ripening) and can greatly influence the ecology of the plant (Purugganan & Fuller, 2009; Meyer *et al*., 2012). The extent to which domestication-induced phenotypic changes affect ecological interactions remains an open question.

It is often proposed that domestication leads to reduced investment in other ecological interactions. In the case of plant-herbivore interactions, a commonly proposed hypothesis suggests that domestication has modified plant traits in ways that decrease resistance to herbivores (Benrey *et al*., 1998). Nonetheless, a meta-analysis (Whitehead *et al*., 2017) conducted in 2017 assessed the overall impact of crop domestication on plant-herbivore interactions and found that, while domestication appears to have a generally negative effect on herbivore resistance, it did not significantly alter plant defense traits. Similarly, a recent study (Glasser *et al*., 2023) exploring the consequences of domestication on plant-pollinator interactions in members of the *Cucurbita* genus revealed that domestication in this group may have promoted greater resource investment in floral traits. This increased investment likely enhances attractiveness to pollinators, potentially increasing reproductive success.

Roots are an important locus for the evolution of non-target traits and species interactions. Root phenotypes have often significantly diverged from their wild relatives due to adaptation to the agricultural environment or as non-target modifications associated with selected traits above ground. Domesticated plants have a different root system architecture, altered root exudate compositions, and significant changes in secondary metabolites across a wide range of species (Meyer *et al*., 2012; Singh & van der Knaap, 2022) which necessarily has an impact in plant interactions with other organisms, including symbiotic microbial communities associated with plant roots (the root or rhizosphere microbiome).

Plants rely on their microbiomes for several functions, including nutrient acquisition, protection against pathogens, and tolerance to biotic stress (Pérez-Jaramillo *et al*., 2018). Often, the root microbiome promotes plant health and growth, contributing to plant development (Singh *et al*., 2020). Given the pivotal importance of root microbiomes for plants and considering the known effects of plant domestication on root systems, in recent years, efforts have been undertaken to decipher the impact of plant domestication on the diversity and assembly of these microbial communities. Despite these efforts, it is still unclear if, or to what extent, plant domestication has modified the ability of the plants to interact with the root microbiome.

Two main hypotheses have been proposed regarding the consequences of domestication: 1) the reduction in microbial diversity resulting from the domestication process and 2) the diminished ability of host plants to shape the assembly of their microbiomes. These hypotheses arise from the idea that the genetic bottlenecks accompanying domestication may lead to the loss of alleles crucial for plant-microbiome interactions, that alterations in root exudates metabolic profiles might further influence microbiome assembly in domesticated plants (Gutierrez & Grillo, 2022), and because of a potential reduction in the benefits of mutualism for domesticated plants. After growing in environments with important nutrient inputs, domesticated plants could reduce the investment in symbiosis, leading to the evolution of reduced symbiont cooperation (Porter & Sachs, 2020). To evaluate these hypotheses, we conducted a meta-analysis comparing studies inoculating domesticated plants and their wild relatives with the same bulk soil in common garden conditions. We found that plant domestication does not result in a uniform reduction in microbial diversity or a consistently diminished ability of host plants to influence their microbiomes. The effects of domestication are species-specific and context-dependent, with some domesticated plants showing increased microbial diversity and structured communities, while others do not. Despite this variation, we found consistent losses and enrichments of microbial taxa in domesticated crops relative to their wild progenitors.

## MATERIALS AND METHODS

### Data collection

To investigate the impact of the domestication process on the diversity and composition of the root microbiome across various crop species, we collected 16S rRNA sequence data from common garden experiments reported in scientific publications. We conducted a comprehensive search in the Scopus and Google Scholar databases using the Boolean search string: Wild [AND] domesticated [AND] microbiome [OR] rhizosphere [AND] crop. Our search centered on contemporary crops with well-identified crop wild relatives, including but not limited to maize, potato, tomato, squash, sorghum, bean, coffee, barley, soybean, cotton, and sunflower. Eligibility for inclusion in our meta-analysis was based on the following selection criteria: 1) the experiments must have been conducted under controlled environmental conditions typical of common garden studies, 2) wild and domesticated variants of the same crop species must have been cultivated concurrently, and inoculated with the same soil, and 3) sequencing must have been performed using Illumina platforms. We excluded studies that assessed the impact of extreme environmental stressors, such as herbivory, but we incorporated those that utilized different soil types (e.g., forest or agricultural soils) as inoculum. Applying these criteria, we narrowed down the pool to 15 publications, representing eight crops: barley, soybean, cotton, tomato, potato, sunflower, maize, and bean. A detailed overview of the number of crop accessions, experimental replicates, and analyzed publications per crop is available in Table S1.

Additionally, due to the large number of papers we could not include in our meta-analysis because of missing raw sequences or metadata, we selected a subset of studies that met our inclusion criteria and reported diversity data and/or ASV tables. A detailed summary of the crop accessions, experimental replicates, and publications analyzed in this secondary dataset is provided in Table S2.

With these datasets, we investigated two primary hypotheses regarding the impact of plant domestication on root microbiomes: 1) the reduction in microbial diversity resulting from the domestication process, and 2) the diminished ability of host plants to control their microbiomes. Since this data did not experimentally assess plant “control”, hereafter we refer as “host influence” to the contribution of plants in shaping root microbiome assemblies. Our analysis focused on the comparison between wild and domesticated categories. To mitigate the potential bias introduced by merging amplicon sequence data from disparate studies, our comparative analyses were restricted to within-study wild and domesticated pairs (W-D pairs). We defined W-D pairs as plants cultivated concurrently and inoculated with the same soil. Typically, each paper contributed a single W-D pair to our analysis. However, in instances where a study investigated more than one soil type, the number of W-D pairs was equivalent to the number of different soil treatments applied.

### Data processing

16S rRNA raw sequences and metadata were downloaded from public repositories, including NCBI, MG-Rast, and Github, and were analyzed with the Quantitative Insights Into Microbial Ecology (QIIME2) platform (Bolyen *et al*., 2019) version 2023.5. For each dataset corresponding to a specific publication, paired-end sequences were analyzed independently. Sequence denoising and quality filtering, including primers and adapters trimming, were performed with the DADA2 plug-in (Callahan *et al*., 2016). Given that all selected studies utilized primers targeting the V4 16S rRNA gene region (e.g., V4, V3-V4, V4-V5), we standardized the analysis by extracting the V4 segment using the 515F–806R primer pairs (Caporaso *et al*., 2011). This approach minimized discrepancies arising from taxonomic assignments across different regions of the 16S rRNA gene. The length of the V4 region sequences was standardized to a range between 120 and 150 nucleotides, based on sequence quality and primer positioning. This criterion was consistently applied across all datasets. Taxonomic classification of the Amplicon Sequence Variants (ASVs) for each dataset was conducted using the “sklearn-base” classifier within QIIME2, referencing the *Greengenes* 2 database (McDonald *et al*., 2024) with a 99% sequence identity threshold. Mitochondrial, chloroplast, and ASVs that were not assignable at the kingdom level were excluded from the analysis. Comprehensive details on the total number of reads (before and after processing) and the count of ASVs for each study have been compiled in Table S3.

### Data Analysis

Data analysis was performed on R software (v.4.3.0) using custom scripts and employing the following packages: Phyloseq (v. 1.46.0) (McMurdie & Holmes, 2013), ggplot2 (v. 3.5.1) (Valero-Mora, 2010), metafor (v. 4.2-0) (Viechtbauer, 2025), vegan (v.2.6-4) (Oksanen, 2017), DESEq2 (v.1.40.2) (Love *et al*., 2014), Source Tracker 2 (Knights *et al*., 2011), and breakaway (v.4.8.4) (Willis & Bunge, 2015). Prior to amplicon analysis, each of the 15 datasets, corresponding to the individual studies included in our meta-analysis, was rarified to a standardized depth of 15,000 reads per sample to ensure comparability across datasets. Specifics on the number of samples and ASVs retained after rarification are available in Table S3. As described previously, all analyses were performed using W-D pairs (pairs of wild and domesticated plants from the same study, species, and treatment).

#### Alpha diversity analyses

To assess the effect of plant domestication on the diversity of the microbiome we calculated Chao1 and Shannon alpha diversity indexes using the *estimate richness* function in the phyloseq package. In addition, we estimated species richness using frequency ratios with the breakaway package. This method applies statistical modeling to predict the number of unobserved taxa, providing flexible modeling of rare species. To test for differences in these indexes we applied the Log Ratio of Means (LogRoM) analysis which compensates for the non-independence of the data by calculating the effect size and variance for each study individually (Friedrich *et al*., 2008). We implemented this test with the *metafor* package, using W-D pairs from all crops and the diversity of the wild plants as reference.

Also, we calculated Rank Abundances Distributions (RADs) for all samples using vegan package, following the methods reported in (Dal Bello *et al*., 2021). From the RAD data, we computed log-linear models to fit linear regressions and compare the slopes between wild and domesticated microbiomes. Absolute slope values were plotted and contrasted; in general, more even communities display smaller slopes.

Additionally, from our secondary dataset (see *Data collection*), we obtained Chao1 and Shannon diversity data either from supplementary information provided in the papers or by downloading ASV tables and using the *estimate richness* function in the phyloseq package. Using these data, we constructed separated databases to test for differences in wild and domesticated microbiomes using the LogRM analysis as previously described.

#### Beta diversity analyses

To explore potential differences in beta diversity between plant categories and crops, we calculated the Bray-Curtis distance for every W-D pair in the dataset. Dissimilarity matrices were analyzed with Permutational Multivariate Analysis of Variance (PERMANOVA) using the *Adonis* function in the vegan package. From the PERMANOVA result, we extracted the R^2^ values to represent the proportion of variation (in terms of beta diversity) explained by category for all W-D pairs and all crops. A dot plot was constructed with this information using ggplot2.

Furthermore, from the dissimilarity matrices we calculated, for all crops, the mean dissimilarity to the bulk of both wild and domesticated plants, and the mean distance to centroid between groups using the *betadisper* function in the vegan package. We plotted both metrics for all W-D pairs in the different crops using ggplot2. Finally, a Bayesian microbial source tracker was performed with the SourceTracker2 program to calculate the contribution of the bulk soil microbial community to the final root microbiome. All samples were rarefied to a depth of 15,000 reads per sample before the analysis.

#### Differentially enriched microbes between categories

We identified differentially enriched microbes between W-D pairs across all crops using the DESeq2 test, implemented in the DESeq2 package. This test was applied exclusively to W-D pairs (excluding bulk soil sequences), with a P-value cutoff of <0.01 and a False Discovery Rate (FDR) correction. Subsequently, we merged the results from all W-D comparisons with differentially abundant features at the family level and selected only features with a log2Fold change of ≤-5 or ≥5, or those present in two or more crops. The results were visualized by crop using forest plots constructed in ggplot2. Additionally, we plotted the individual abundances of microbial families enriched in the same category across multiple crop species (Fig. S6).

## RESULTS

We conducted a systematic meta-analysis to assess the influence of plant domestication on root microbiomes across different plant species. After applying our inclusion criteria and filters of data quality, 15 independent studies were retained to perform the full analyses. These studies encompass eight different crop species and included 947 individual plants belonging to maize (n = 128), wheat (n= 172), barley (n= 55), sunflower (n= 127), tomato (n= 169), cotton (n= 31), bean (n= 52), and soybean (n= 160) (Table S1). Except for soybean, and sunflower, all crops were represented by more than one accession of wild and domesticated plants (Table S1).

Additionally, we included diversity metrics from studies that met our inclusion criteria but were excluded from the full analyses due to missing data. This secondary dataset included eight independent studies from six different crop species, including 758 individual plants belonging to: tomato (n= 28), potato (n= 210), maize (n= 264), sorghum (n= 100), rice (n= 115), and agave (n= 41) (Table S2). All crops in this dataset were represented by more than one accession of wild and domesticated plants, increasing the host genetic diversity analyzed in this study.

In total, we analyzed 40 different W-D pairs (pairs of wild and domesticated plants from the same study, species, and treatment). This included 23 W-D pairs from the main dataset, used for the full analyses, and 17 W-D pairs from the secondary dataset, used exclusively for diversity comparisons.

### Domesticated plants show increased microbiome diversity

To test for differences in microbial diversity between wild and domesticated plants we performed multiple diversity, richness, and evenness analyses. Contrary to the common expectation that domestication reduces microbiome diversity, we found, on average, domesticated plants exhibited slightly higher diversity in terms of Chao1 index and species richness calculated using frequency ratios. This pattern was consistent across both the main dataset that included the raw sequences (LogRoM: 0.01, 95% CI: −0.06-0.7, *p* < 0.000, Fig. 1 - Chao1; LogRoM: 0.01, 95% CI: −0.01-0.8, *p* < 0.000, Fig. S1 - Frequency ratios) and the secondary dataset (LogRoM: 0.04, 95% CI: −0.08-0.17, *p* < 0.000, Fig. S3b - Chao1 index). Despite these differences in estimated richness, wild and domesticated species tended to have similar diversity using the Shannon index (main dataset: LogRoM: −0.01, 95% CI: −0.01-0.0, *p*= 0.1, Fig. S2; secondary dataset: LogRoM: 0.00, 95% CI: −0.02-0.02, *p*= 0.1 (without agave data - figure not shown)). The main exception to this trend was the comparison using the Shannon index for Agave samples in the secondary dataset, where we found higher diversity in wild plants, influencing the overall model toward greater diversity in wild plants (LogRoM: −0.01, 95% CI: −0.02-0.01, *p* < 0.00, Fig. S3a).

**Figure 1.**
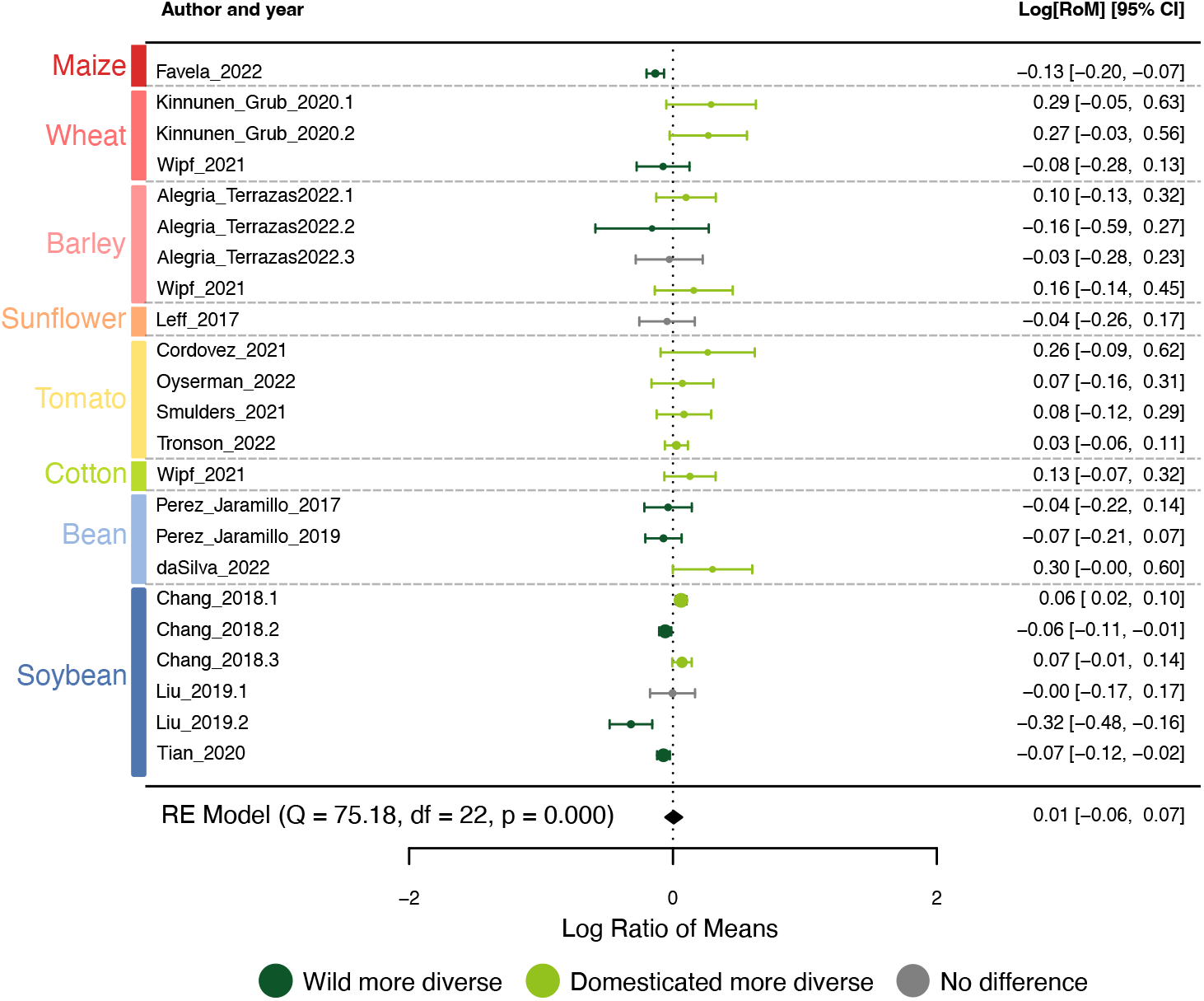
Effect of Domestication on Microbial Alpha Diversity Across Different Crop Species. Forest plot showing the log Ratio of Means (LogRoM) using Chao1 diversity index for all 23 Wild-Domesticated (W-D) pairs. The mean microbial diversity of wild plants was used as reference. Dashed lines denote the separation between different plant species. Bars represent the mean response ratios with 95% confidence intervals. Colors indicate the results: dark green represents higher diversity in wild plants, light green represents higher diversity in domesticated plants, and grey indicates no statistical difference.

Most pairwise comparisons (pairs of wild and domesticated plants from the same study, species, and treatment) showed differences in diversity, but the effects were highly variable across species (Fig.1). Among the 23 Wild-Domesticated (W-D) pairs evaluated, only three (13%) exhibited no statistical difference. Of the 19 (82%) pairs that showed statistically significant differences, 12 (63%) displayed increased diversity in domesticated plants, whereas eight (42%) had higher diversity in their wild counterparts. Whereas for the secondary dataset (Fig. S3b), we found that 77% of the W-D pairs analyzed showed significant differences, from those, 71% showed increased diversity in domesticated plants, while 29% showed higher diversity in wild plants. Notably, the effect of domestication appears to be species-specific and related to functional groups. Crops like wheat, tomato, and cotton showed increased diversity in their domesticated forms compared to their wild forms, whereas maize and legumes (beans and soybeans) presented greater diversity in their wild forms. Despite these overall patterns, we observed different outcomes, even within the same species. The only exception to this inconsistency was observed in tomatoes, where all four comparisons consistently revealed higher diversity in domesticated plants. The most variable crop species was soybean, where of the six total W-D pairs analyzed, two resulted in higher diversity in domesticated versions, three in higher diversity in wild plants, and one showed no statistically significant difference. Furthermore, we did not find differences in microbiome evenness between wild and domesticated plants; when comparing absolute slope values calculated from the RADs data, we found that all analyzed microbial communities had small slopes, indicating highly even communities regardless of domestication category (Fig. S4).

### Host species effect is more important than domestication in shaping microbiome composition

We then evaluated the effect of domestication on microbiome composition using PERMANOVA. We found that the domestication category accounted for a relatively small fraction of the variation across most of the crop species studied (Fig. 2 and Fig. S5), suggesting a potential species-specific effect. For several species, including maize, wheat, barley, sunflower, tomato, and bean, domestication explained approximately 10% of the observed variance in microbial community composition. Notably, cotton and soybean (except for two pairs) stood out, with domestication accounting for a larger portion of the variance, ranging from 12% to 25%. Additionally, we leveraged experiments that used multiple soil types to inoculate plants, allowing us to test the contribution of the inoculum to the final microbial community by separating wild and domesticated plants independently (Table S4). Although only a small set of papers had different soil types, for barley, different initial soils accounted for a minor portion of the total variation (9.8% for wild plants and 10.9% for domesticated plants). In contrast, for soybeans, using soils from different origins accounted for a significant percentage of the variation (~75% for wild and ~78% for domesticated plants). Taken together, these results suggested that domestication could affect the degree of host influence but that there might be variation across species.

**Figure 2.**
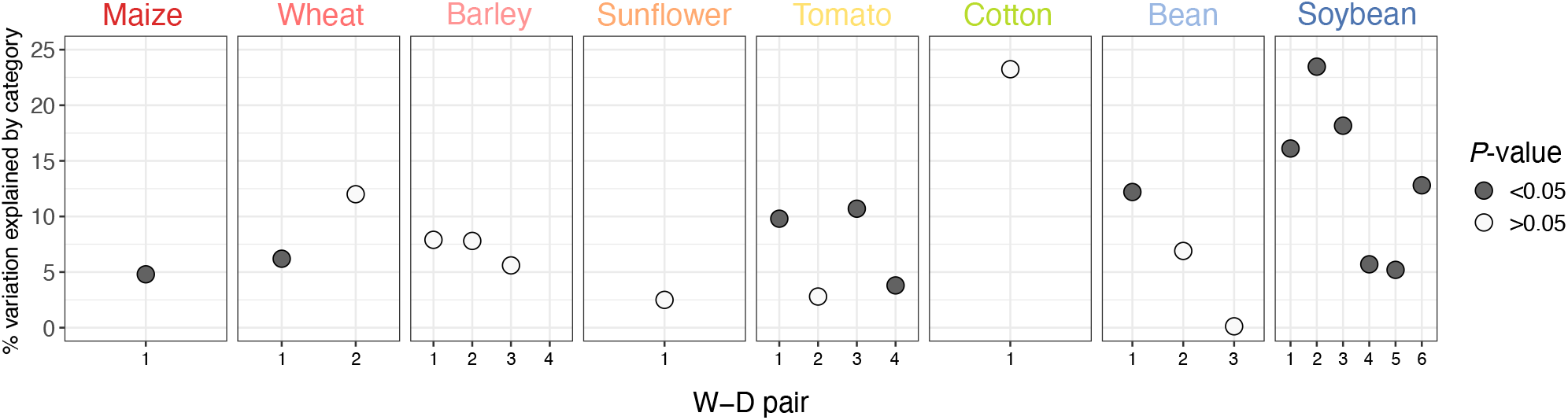
Effect of Domestication on Microbial Community Composition Across Different Crop Species. Dot plot showing the percentage of variation in microbial community composition explained by domestication category (wild vs. domesticated), according to the Bray-Curtis distance measure. Numbers in X axis represent the number of Wild-Domesticated pairs (pairs of wild and domesticated plants from the same study, species, and treatment) analyzed per crop.

### Wild relatives do not display systematic evidence of increased host influence of their microbiomes

To evaluate the extent of host influence exerted by wild and domesticated plants over their microbiomes, we leveraged beta diversity metrics, comparing the microbial communities associated with plants to those from bulk soils, which served as a reference for the initial microbial community (Fig. 3). We hypothesized that if plants have a relatively strong influence in the assembly of their microbiomes, then we would expect to find: a) significant dissimilarity between plant-associated microbiomes and bulk soil microbiomes (Fig. 3a), suggesting that plants are selecting specific microbial assemblages distinct from the initial soil; b) low intra-group dispersion (Fig. 3b), indicating that plant replicates of the same genotype consistently select for similar microbial compositions; and c) less proportion of the final community explained by the initial bulk soil (Fig. 3c), implying that the host plays a decisive role in shaping its microbiome, potentially steering it away from the original soil community composition.

**Figure 3.**
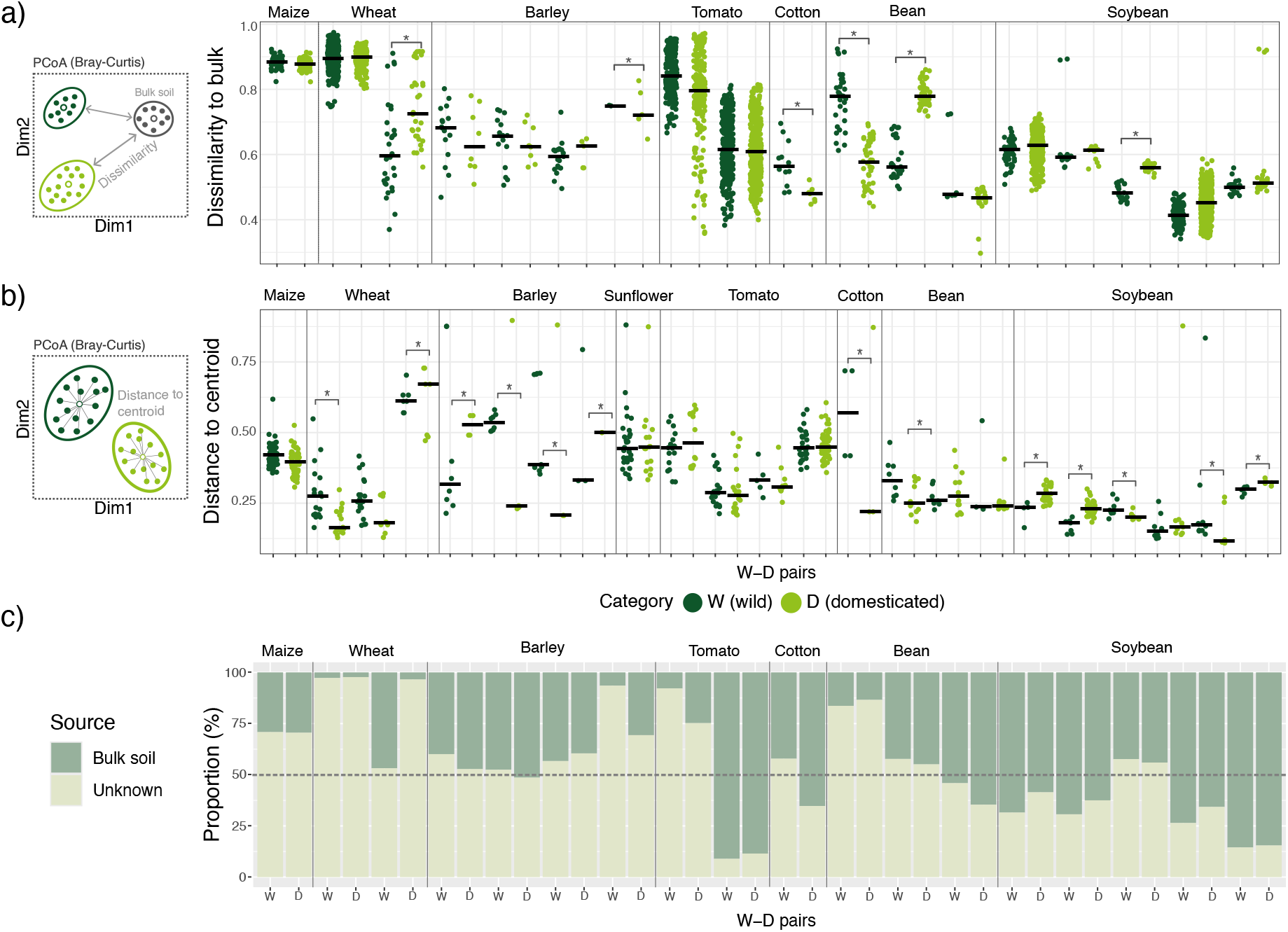
Impact of Domestication on Host Influence of the Microbiome. **A)** Bray-Curtis dissimilarity to bulk soil for wild-domesticated (W-D) pairs across different crop species. Statistically significant differences are indicated by an asterisk (Wilcoxon rank-sum test, p < 0.05). This analysis includes only experiments that included bulk soil sequences. **B)** Intra-group dispersion for all W-D pairs analyzed using Bray-Curtis distance. Statistically significant differences are indicated by an asterisk (Wilcoxon rank-sum test, p < 0.05). **C)** Source Tracker analysis estimating the contribution of bulk soil to the final microbial community of wild and domesticated plants across different species.

Overall, we did not identify a consistent pattern linking host influence to the category of domestication. Most comparisons were not statistically significant. Among the significant comparisons, we observed greater dissimilarity to bulk in wild barley and cotton and in domesticated wheat and soybean (Fig. 3a). Similarly, we found a diverse range of outcomes in the intra-group dispersion analyses (Fig. 3b). Domesticated plants generally exhibited more intra-group dispersion in barley and soybean, whereas we found the opposite effect (more intra-group dispersion in wild varieties) in wheat, cotton, and bean. Maize, sunflower, and tomato showed no differences. We didn’t find a clear pattern in the contribution of bulk soil microbiomes to the final plant-associated communities across different levels of domestication (Fig. 3c).

We explored if some other plant attribute could better account for our results. Grouping our results by functional plant group rather than domestication status provided increased insight. Grasses (maize, wheat, barley) consistently demonstrated a more distinct divergence from their initial bulk soil microbiomes (Fig. 3a), with less than 50% of the final community composition attributable to the bulk soil (Fig. 3c). Whereas the legume soybean presented final communities closely aligned with the initial bulk soil, reflecting less than 50% differentiation.

### Some microbial taxa differentially associate with plants across species according to the domestication status

Finally, we investigated if, despite all the variation, we could identify microbial groups differentially present in wild and domesticated crops (see methods; Fig. 4). We identified a few families consistently enriched or depleted from domesticated crops (relative to their wild counterparts) across different crops. Two members of the *Rhizobiaceae* family (*Rhizobiaceae_A_500471* and *Rhizobiaceae_A_500279*) were significantly enriched in the domesticated variants of wheat, tomato, and soybean. Whereas two members of the *Burkholderiaceae* family (*Burkholderiaceae_A_592522* and *Burkholderiaceae_A_574758)* were consistently enriched in the wild counterparts of wheat, tomato, bean, and soybean. We found these *Burkholderiaceae* taxa in both the domesticated and the wild rhizosphere. In contrast, the *Rhizobiaceae* members appeared almost absent in the wild versions of wheat and tomato (minimum absolute abundance < 50 reads; Fig. S6).

**Figure 4.**
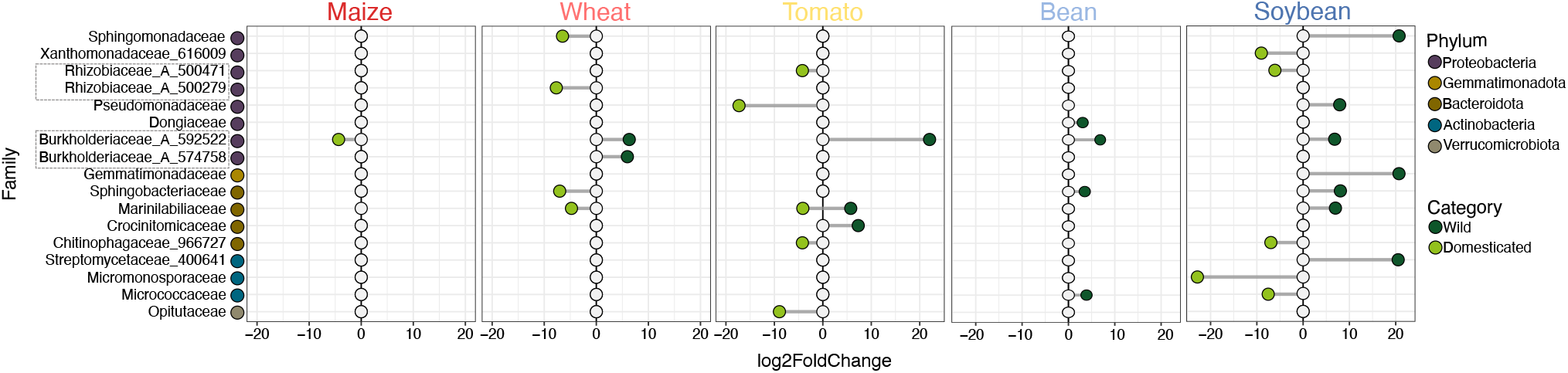
Differentially enriched microbial families between wild and domesticated plants across various crop species. DESeq analysis at the family level displaying microbes either enriched in two or more crop species or exhibiting a fold change greater than +5 or less than −5.

## DISCUSSION

Our study provides new insights into the relationship between plant domestication and microbial diversity and assembly, challenging the widely held belief that domestication always leads to a decrease in microbial diversity and impairs the plant’s influence in the assembly of its microbiome. Our results indicate that while domestication does have an effect on microbial diversity and community composition, these effects vary widely across species and experimental conditions.

Contrary to the hypothesis that domestication generally reduces microbial diversity, we found a small but significant reduction of microbiome diversity in the rhizosphere of wild relatives (Fig. 1, Log Ratio of Means with Chao1 index, *p* < 0.001). In fact, for the Chao1 index as well as for the frequency ratios test, most of the significant comparisons indicated that domesticated plants exhibited higher microbial diversity than wild plants (Fig. 1 and Fig. S1). When using the Shannon diversity index, we observed fewer differences and mostly saw no change in diversity across comparisons. The hypothesis that domestication leads to a reduction in microbiome diversity is largely based on the idea that domestication reduces genetic diversity, potentially leading to the loss of alleles important for plant-microbe interactions through selective bottlenecks (Gutierrez & Grillo, 2022) and that in domesticated plants, a reduced investment in symbiosis could promote the evolution of reduced symbiont cooperation (Porter & Sachs, 2020).

It is not entirely clear why the reduction of genetic diversity at the level of populations would lead to reduced microbiome diversity at the level of individuals. Instead, we might expect a decrease in microbiome diversity across hosts (less dispersion) in domesticated varieties compared to their wild counterparts. However, our results highlight that this is only the case in some pairs –mostly wheat, barley, and cotton. The opposite is true in some other pairs, including many of the soybean comparisons (Fig. 3). One of the limitations in making clear conclusions about the impact of host diversity on the microbiome is that host diversity is rarely controlled in these studies (for example, many domesticated accessions might be compared against a single population of wild relatives).

Even the loss of alleles important for plant-microbe interactions does not have to be associated with decreased microbiome diversity. It has been proposed, for example, that a reduction in genetic diversity might lead to less efficient microbiome filtering mechanisms, allowing domesticated plants to be less selective and thus associate with a greater diversity of microbes (Soldan *et al*., 2021; Favela *et al*., 2022; Gutierrez & Grillo, 2022). This hypothesis has gained attention due to its potential implications for agriculture and resilience to climate change. The variation in microbial communities explained by host genotype represents the substrate upon which natural selection can act and, therefore, it is also the variation that could be manipulated for agricultural improvement (Brown *et al*., 2020). Thus, a reduction in host influence resulting from domestication could lead to decreased evolutionary potential within microbial communities, potentially limiting the adaptive resources of plants in response to environmental stresses (Soldan *et al*., 2021). This hypothesis of less efficient filtering mechanisms in domesticated crops is consistent with our alpha (within-plant) diversity results, where we observed a small increase in microbiome diversity in domesticated varieties. But, contrary to this expectation, we found little support for decreased host influence in domesticated plants. In most pairs, the microbial community of the rhizosphere is equally distant to the bulk soil, with soybean being the only consistent example of increased distance to bulk soil in the domesticated varieties (even if most are not significant) (Fig. 3).

Until recently (Doebley *et al*., 2006), the existence of a domestication syndrome had promoted the idea that most plant species that undergo domestication suffer the same phenotypic consequences. Likewise, this could have also promoted the assumption that, for all crop species, domestication would always negatively affect microbial diversity. In addition to finding, on average, increased microbial diversity in domesticated plants, our analysis revealed significant variability inside species. For many of the crops analyzed, there were instances where wild plants exhibited higher microbial diversity, while in other cases, domesticated plants showed greater diversity. We found this variability in different studies using the same plant accessions and even within the same study when different initial soils were used, implying a significant role of genotype-by-environment (GxE) interactions in shaping microbiome assembly.

It is known that microbial communities from different habitats (i.e., different initial soils) can exhibit distinct responses to the same host genotype. It has been proposed that environmental heterogeneity can decouple host genotypes from their microbiomes by altering genotype-microbe relationships (Bradshaw, 1965; Wagner *et al*., 2016), potentially promoting mixed results even when controlling for host genotypes. Our results suggest that domestication’s effects on microbial diversity are not universally negative and may vary depending on the specific crop species and environmental conditions involved. This observation is further supported when examining the impact of domestication on microbial community composition. Although most comparisons showed differences in beta diversity (Figure S5), domestication status accounted for a relatively minor fraction of the microbial variation across the analyzed species (Fig. 2). Instead, much of the variation, at least in studies using more than one soil type, is explained by the soil origin (Table S4). This finding aligns with the results of individual studies on rice (Edwards *et al*., 2015), poplar (Bonito *et al*., 2019), soybean (Liu *et al*., 2019), and the legume *Medicago truncatula* (Brown *et al*., 2020), where soil origin is a stronger predictor of community composition than host genotype. Some of these studies suggest that endophytes are more sensitive to host genotype signals, while rhizosphere communities are more affected by the environment (Brown *et al*., 2020). Future studies looking at the effect of domestication should aim to separate the effects of domestication on different microbial functional groups.

Despite all the variation found between crops, we detected two members of the *Rhizobiaceae* family consistently enriched in domesticated wheat, tomato, and soybean, and two members of the *Burkholderiaceae* family enriched in wild wheat, tomato, bean, and soybean (Fig. 4). In some crop species, domesticated plants exhibit increased photosynthetic capacity compared to wild relatives, which is thought to support the development of larger grains or fruits (Koester *et al*., 2016; Brestic *et al*., 2018). This greater carbon fixation could be increasing the root exudation in domesticated plants, potentially attracting more rhizobia and other copiotrophs in the rhizosphere (Richards, 2000). Additionally, prolonged nitrogen fertilization in agricultural systems is proposed to promote the evolution of less-cooperative rhizobia in domesticated cultivars, potentially leading to higher root abundance of *Rhizobiaceae* members, many of which may not contribute significantly to plant nutrition (Weese *et al*., 2015; Liu *et al*., 2019). Particularly, rhizobia with poor or null symbiotic effectiveness are more prevalent in agricultural soils and are predominantly associated with cultivated crops (Sachs *et al*., 2010). Thus, a high abundance of those microbes does not necessarily indicate a higher nitrogen fixation in the rhizosphere of domesticated crops.

On the other hand, the consistent enrichment of *Burkholderiaceae* family members in the wild versions of multiple species aligns with previous research on common beans and rice (Pérez-Jaramillo *et al*., 2019; Wu *et al*., 2022). Although the mechanisms behind wild plants harboring an increased abundance of those particular microbes remain unknown, it has been shown that members of the *Burkholderiaceae* family are known for their ability to promote plant growth and suppress soil-borne pathogens, supporting plant health overall (Estrada-De Los Santos *et al*., 2001). One potential explanation is that compared to domesticated plants, wild relatives produce a greater diversity of secondary metabolites, which may promote the recruitment of these beneficial microbes (Ku *et al*., 2020). Nonetheless, for the case of the common bean, positive associations with *Burkholderiaceae* family members were found only when using native soils (e.g. versus agricultural soils) (Pérez-Jaramillo *et al*., 2019), suggesting potential local adaptation to microbes found within the natural distribution of such species.

## CONCLUSIONS AND FUTURE DIRECTIONS

In this study, we challenged accepted hypotheses regarding the consequences of plant domestication on root microbiomes. Contrary to the expectation that domestication generally reduces microbial diversity, we found a small but significant reduction of microbiome diversity in the rhizosphere of wild relatives. Furthermore, by analyzing multiple crop species, we did not find sufficient evidence to support the hypothesis that domestication negatively affects host influence on the assembly of the microbiome. The consistent enrichment of certain bacterial families in both wild or domesticated versions of different crop species (*Rhizobiaceae* for domesticated and *Burkholderiaceae* for wild plants) could be suggesting, at least for the data analyzed in this work, that the effects of domestication may be stronger at the level of recruiting specific microbes rather than affecting the entire community. Overall, our results suggest that the impacts of domestication on the root microbiomes are not universally negative in terms of microbial diversity. Instead, they appear to be species-specific and contingent upon environmental factors.

Despite our findings, much work remains to fully understand the ecological and evolutionary consequences of plant domestication on root microbiomes. First, diversity is not everything; characterizing functional traits of microbiomes associated with wild and domesticated plants is crucial to better understanding the consequences of domestication for plant-microbe interactions. Second, it is possible that the effect of domestication would be more pronounced with close symbionts and endophytes, more work is necessary to understand how domestication can alter these interactions. Third, it is well-documented that domestication significantly impacts root system architecture, but there is limited research on how such changes affect the microbes inhabiting the roots. How changes in root size, phenotype, and physical properties of the roots impact the recruitment of microbes remains as open questions. Similarly, although exudates have been studied more than the physical structure of roots for microbiome recruitment, this is still an area where more research is needed to identify general patterns.

Critical directions for future research are to incorporate plant diversity in the design or analysis of the experiments to start uncovering the connection between genetic changes of domesticated crops and their impact on the microbiome. In particular, controlling for genetic diversity between plants is important to be able to attribute differences between groups to the process of domestication itself. Similarly, to evaluate the role of domestication on the loss of host influence, it will be important to incorporate native soils into common garden experiments. Local adaptation is important for wild populations, and relying exclusively on agricultural soils (or other soils outside the natural range of wild plants) may hinder significant microbial interactions that occur only with the specific set of microbes to which wild plants are naturally adapted.

Finally, introgression between wild and domesticated plants is a common phenomenon, especially at the centers of origin of the species (Ellstrand *et al*., 2002). The evolutionary consequences of this have been extensively studied from a genetic perspective (Janzen *et al*., 2019), and as a source for key agronomic traits (Le Corre *et al*., 2020). Nonetheless, even though there is increasing evidence showing that hybridization impacts root microbiomes (Bouffaud *et al*., 2014; Zhang *et al*., 2023), the consequences of introgression between wild and domesticated plants remain unclear. Recently, studies on rice (Zhang *et al*., 2023) and maize (Wagner *et al*., 2021) have found that both root and seed microbiomes could contribute to heterosis (“hybrid vigor”) in hybrids, providing evidence of the important role of microbiomes in the fitness of inbreds. Studying the potential adaptive and maladaptive consequences of microbiome introgression is a new and exciting area of study, with significant implications for plant breeding and conservation biology.

## ACKNOWLEDGMENTS AND FUNDING SOURCES

We would like to thank Brandon S. Gaut, Eduardo Choreño-Parra, Santiago Rosas-Plaza, Celia Symons, and members of the COMMONS Lab for productive discussions about this work. AHT was supported by the Human Frontiers Science Program Long-Term Postdoctoral Fellowship (LT 0047/2023-L) and the University of California President’s Postdoctoral Fellowship. AEH was supported by UNAM-PAPIIT: IN214124. MRG was supported by the School of Biological Sciences at the University of California, Irvine (UCI).

## AUTHOR CONTRIBUTIONS

AHT and MRG designed the research with input from AE. AHT performed all statistical analyses. AHT wrote the manuscript with substantial feedback from MRG and AE.

## DATA AVAILABILITY STATEMENT

Data and R code for analyses is available on Github at: https://github.com/AlejandraHT/Plant_domestication_microbiomes

