## Supplementary information for "Plant domestication does not reduce diversity in root microbiomes"

**This PDF file includes:**

- Figures S1 to S6
- Tables S1 to S4
- List of full references of scientific papers used in the meta-analysis

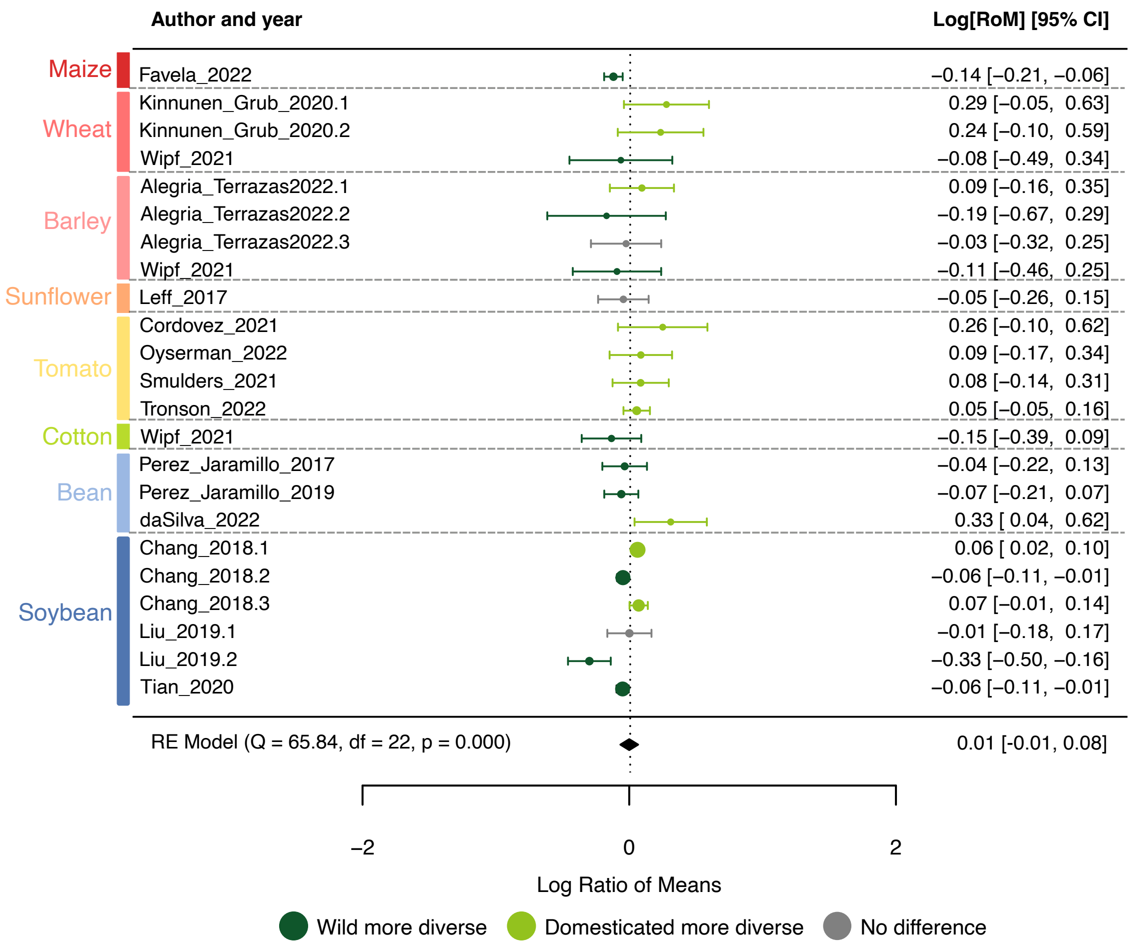


**Fig. S1.** Forest plot showing the log Ratio of Means (LogRoM) using frequency ratios of species richness for all 23 Wild-Domesticated (W-D) pairs from the main dataset. The mean microbial diversity of wild plants was used as reference. Bars represent the mean response ratios with 95% confidence intervals.

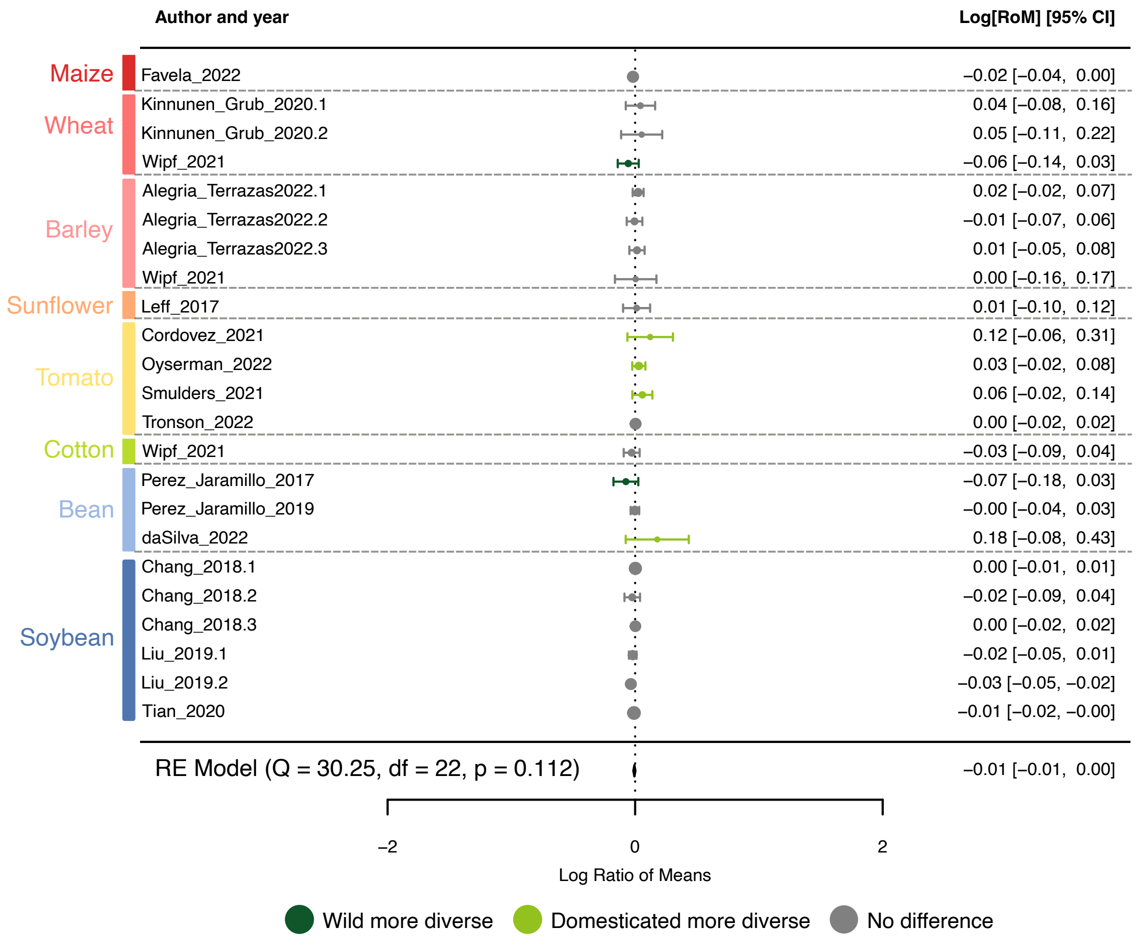


**Fig. S2.** Forest plot showing the log Ratio of Means (LogRoM) using Shannon diversity index for all 23 Wild-Domesticated (W-D) pairs from the main dataset. The mean microbial diversity of wild plants was used as reference. Bars represent the mean response ratios with 95% confidence intervals.

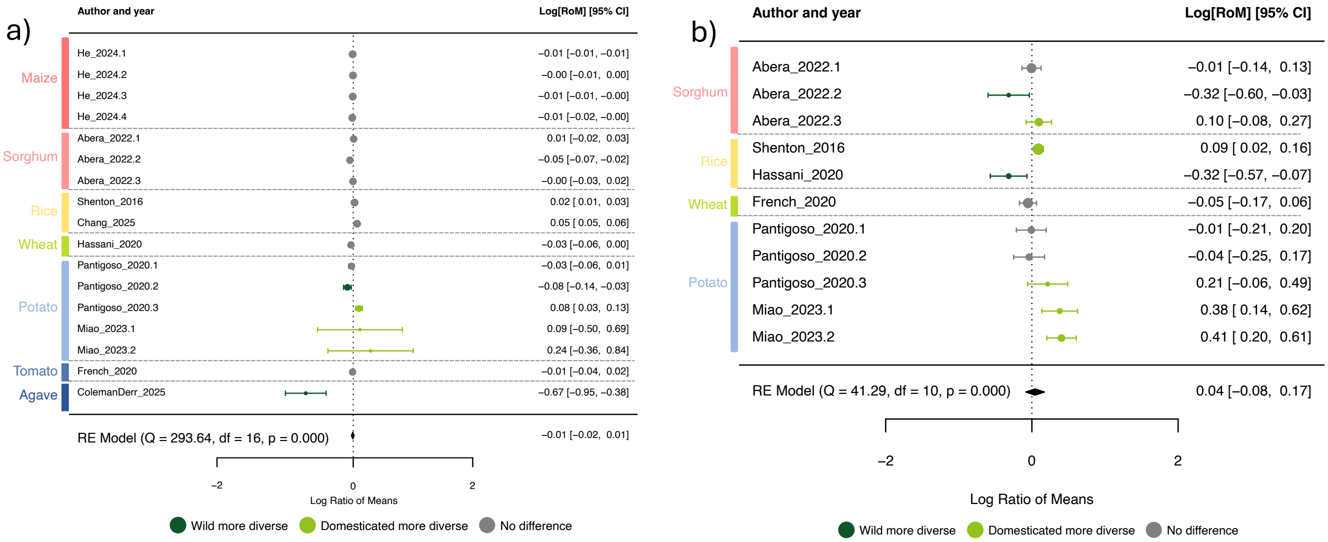


**Fig. S3. Forest plot showing the log Ratio of Means (LogRoM) of diversity indexes for all Wild-Domesticated pairs from the secondary dataset.** a) LogRoM using Shannon index. b) LogRoM using Chao1 index. For both, the mean microbial diversity of wild plants was used as reference. Bars represent the mean response ratios with 95% confidence intervals. Fewer W-D pairs were analyzed in terms of richness (Chao1 – b)) due to missing ASV tables.

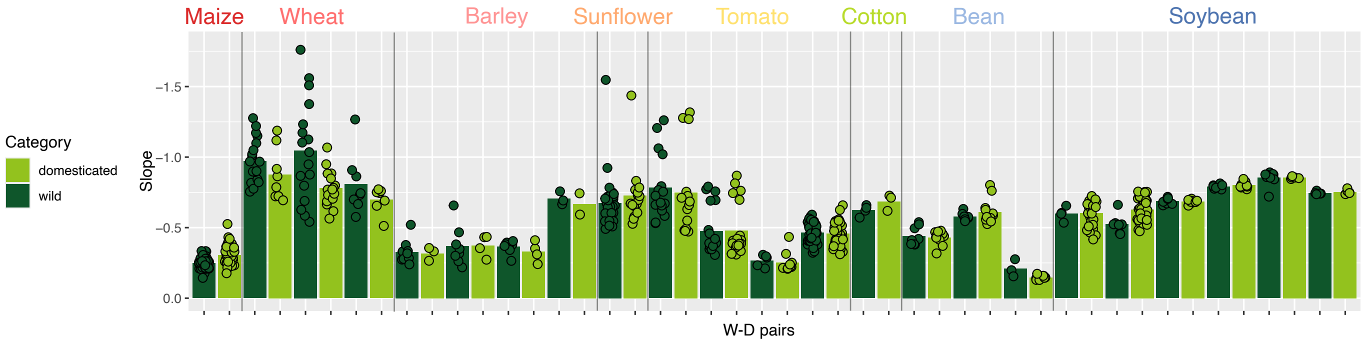


**Fig. S4. Average absolute values of slopes calculated from log-linear Rank Abundance Distributions (RADs) for each W-D pair.** Values were calculated by fitting linear regressions on RADs for all samples. More even communities display smaller slopes.

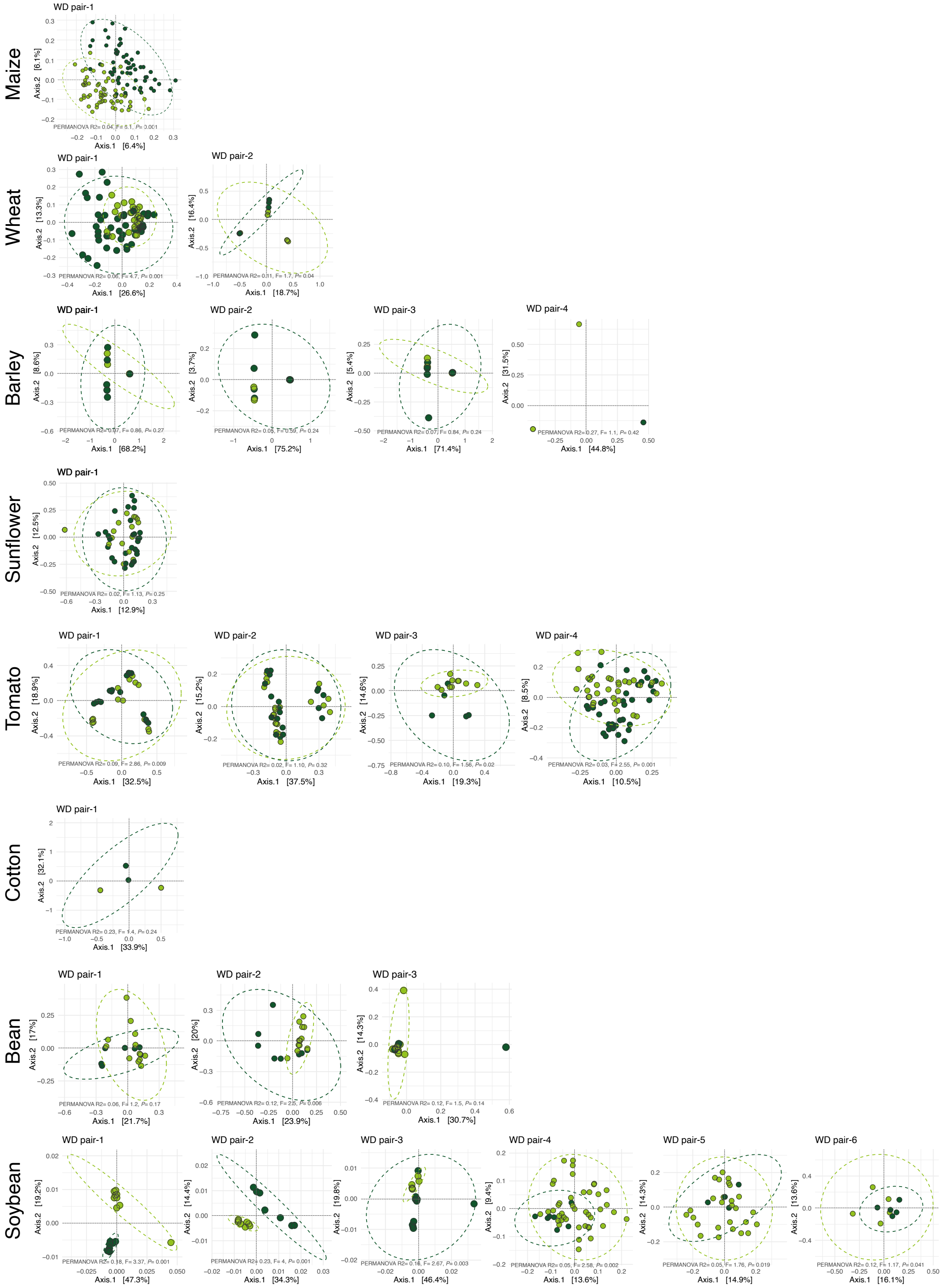


**Fig. S5. Principal Coordinate Analysis (PCoA) with Bray-Curtis distance and PERMANOVA result from all W-D pairs analyzed.** For all plots, Wild = dark green, Domesticated = light green.

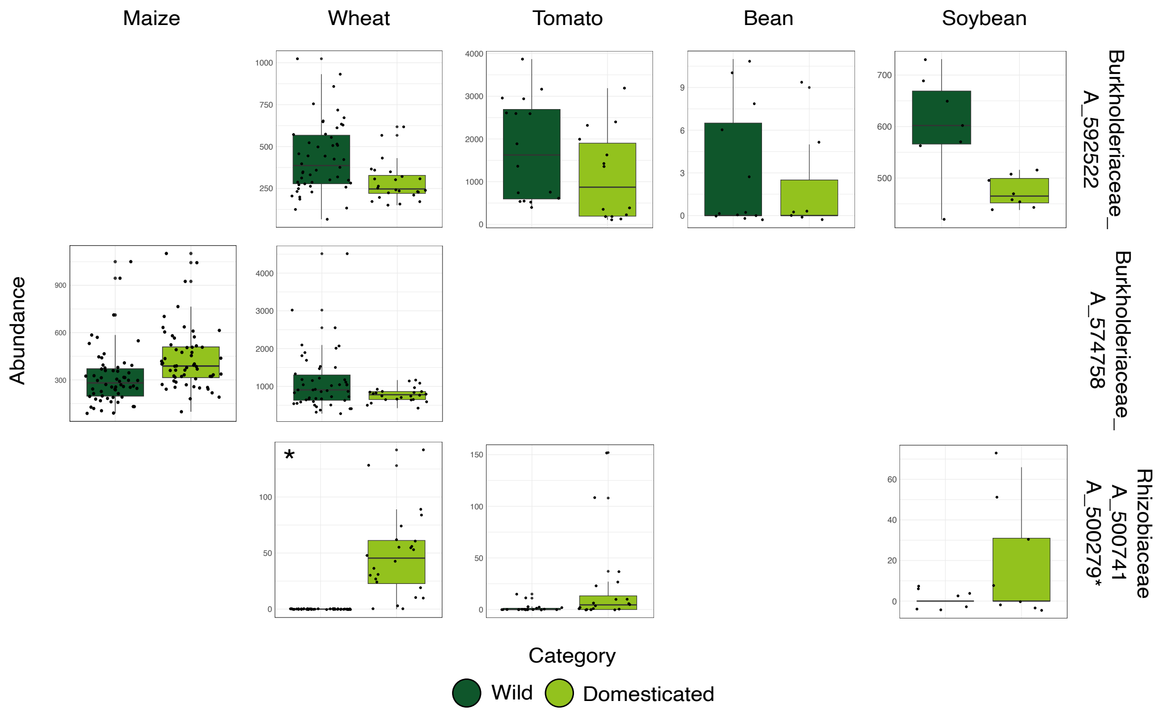


**Fig. S6. Absolute abundance per plant category of microbial groups found significant in DESEQ2 analysis.**

**Table S1. Total number of plants replicates and accessions analyzed per crop in the main dataset**.
*# of plants* refers to the total number of individuals analyzed per crop. *# of plant accessions* refer to the number of genetically distinct individuals used per crop.

| Crop | # wild plants | # wild plant accessions | # dom plants | # dom plant accessions | # total replicates | # papers | Reference |
|---|---|---|---|---|---|---|---|
| **Maize** | 65 | 6 | 63 | 6 | 128 | 1 | (Favela *et al.*, 2022) |
| **Wheat** | 94 | 6 | 78 | 4 | 172 | 2 | (Kinnunen-Grubb *et al.*, 2020; Wipf & Coleman-Derr, 2021) |
| **Barley** | 35 | 3 | 20 | 2 | 55 | 2 | (Wipf & Coleman-Derr, 2021; Alegria Terrazas *et al.*, 2022) |
| **Sunflower** | 64 | 1 | 63 | 1 | 127 | 1 | (Leff *et al.*, 2017) |
| **Tomato** | 82 | 3 | 87 | 2 | 169 | 4 | (Cordovez *et al.*, 2021; Smulders *et al.*, 2021; Oyserman *et al.*, 2022; Tronson *et al.*, 2022) |
| **Cotton** | 21 | 2 | 10 | 1 | 31 | 1 | (Wipf & Coleman-Derr, 2021) |
| **Bean** | 18 | 2 | 34 | 4 | 52 | 3 | (Pérez-Jaramillo *et al.*, 2017, 2019; da Silva *et al.*, 2023) |
| **Soybean** | 44 | 1 | 116 | 1 | 160 | 3 | (Chang *et al.*, 2019; Liu *et al.*, 2019; Tian *et al.*, 2020) |

**Table S2. Total number of plants replicates and accessions analyzed per crop in the secondary dataset.** # of plants refers to the total number of individuals analyzed per crop. # of plant accessions refer to the number of genetically distinct individuals used per crop.

| Crop | # wild plants | # wild plant accessions | # dom plants | # dom plant accessions | # total replicates | # papers | Reference |
|---|---|---|---|---|---|---|---|
| **Tomato** | 7 | 2 | 21 | 6 | 28 | 1 | (French *et al.*, 2020) |
| **Potato** | 88 | 6 | 122 | 8 | 210 | 2 | (Pantigoso *et al.*, 2020; Miao & Lankau, 2023) |
| **Maize** | 132 | 11 | 132 | 7 | 264 | 1 | (He *et al.*, 2024) |
| **Sorghum** | 20 | 2 | 80 | 8 | 100 | 1 | (Abera *et al.*, 2022) |
| **Rice** | 62 | 17 | 53 | 13 | 115 | 2 | (Shenton *et al.*, 2016; Chang *et al.*, 2025) |
| **Agave** | 29 | 2 | 12 | 1 | 41 | 1 | (Coleman-Derr *et al.*, 2016) |

**Table S3. Number of sequences retained after filtering process and total Amplicon Sequence Variants (ASV) analyzed per paper.** In *total reads by category*: W= wild plants, D= domesticated plants.

| Author/Year | Crop | Raw reads | Reads after filtering | Total reads by category | Total ASVs | ASVs after rarefaction |
|---|---|---|---|---|---|---|
| **Alegria-Terrazas et al, 2022** | Barley | 9,432,335<br>Min: 36,212<br>Max:892,576 | 5,500,276<br>Min:21,274<br>Max:472,491 | W= 2,760,693<br>D=1,011,187 | 20,159 | 10,415 |
| **Chang et al, 2018** | Soybean | 4,558,947<br>Min:79,604<br>Max:80,410 | 1,010,436<br>Min:15,369<br>Max:21,077 | W=444,190<br>D=419,806 | 3,188 | 2,848 |
| **Cordovez et al, 2021** | Tomato | 1,851,413<br>Min:23,682<br>Max:65,595 | 1,103,720<br>Min:14,525<br>Max:41,433 | W=401,959<br>D=394,089 | 3,912 | 3,825 |
| **Favela et al, 2021** | Maize | 14,774,081<br>Min:<br>Max:410,789 | 10,504,418<br>Min:957<br>Max:293,909 | W=5,711,684<br>D=4,702,635 | 38,309 | 34,074 |
| **Kinnunen-Grub et al, 2020** | Wheat | 24,215,139<br>Min:1701<br>Max:289,546 | 5,784,498<br>Min:2<br>Max:175,905 | W=1,713,823<br>D=1,161,633 | 4,402 | 2,317 |
| **Leff et al, 2017** | Sunflower | 14,783,123<br>Min:76<br>Max:117,868 | 10,682,174<br>Min:11<br>Max:105,374 | W= 610,461<br>D= 360,710 | 26,098 | 9472 |
| **Liu et al, 2018** | Soybean | 19,358,039<br>Min:40,987<br>Max:412,663 | 4,862,423<br>Min:4,911<br>Max:152,583 | W=556,435<br>D=2,347,034 | 10,421 | 6,547 |
| **Oyserman et al, 2022** | Tomato | 61,652,169<br>Min:501<br>Max:570,931 | 31,000,982<br>Min:10<br>Max:345,787 | W=1,072,406<br>D=1,272,152 | 13,707 | 11,234 |
| **Perez-Jaramillo et al, 2017** | Bean | 4,047,273<br>Min:91,945<br>Max:131,648 | 2,165,082<br>Min:39,579<br>Max:72,884 | W=489,319<br>D=770,461 | 6,023 | 5,623 |
| **Perez-Jaramillo et al, 2019** | Common bean | 3,119,178<br>Min:86,537<br>Max:119,210 | 1,090,726<br>Min:19,157<br>Max:42,910 | W=264,168<br>D=423,188 | 2,718 | 2,629 |
| **Smulders et al, 2021** | Tomato | 7,271,546<br>Min:122,787<br>Max:330,479 | 2,788,200<br>Min:40,444<br>Max:130,352 | W=492,545<br>D=971,158 | 10,027 | 8,484 |
| **Tian et al, 2020** | Soybean | 1,600,222 | 285,630 | W=76,130 | 1,388 | 1,374 |

| | | | | | | |
|---|---|---|---|---|---|---|
| | | Min: 79,552<br>Max: 80,397 | Min: 13,036<br>Max: 15,394 | D=67,251 | | |
| **Tronson et al, 2020** | Tomato | 9,868,547<br>Min:16,882<br>Max:73,052 | 5,230,298<br>Min:303<br>Max:42,877 | W=919,752<br>D=938,949 | 19,334 | 10,662 |
| **Wipf et al, 2021** | Barley/<br>Wheat/<br>Cotton | 7,614,252<br>Min:296<br>Max:180,707 | 4,431,837<br>Min:0<br>Max:134,450 | Wheat<br>W=234,021<br>D=299,945<br>Cotton<br>W=190,419<br>D=117,414<br>Barley<br>W=134,153<br>D=91,908 | 15,478 | 5,769 |
| **DaSilva et al, 2022** | Lima bean | 2,734,632<br>Min: 97,425<br>Max: 233,060 | 2,462,741<br>Min:86,344<br>Max:210,279 | W=356,955<br>D=1,433,061 | 11,072 | 8,648 |

**Table S4. PERMANOVA results from experiments using multiple soil types as inocula.**

| | | PERMANOVA result | | |
|---|---|---|---|---|
| **Crop** | **Category** | **R2 * 100** | **F** | ***P* value** |
| Barley | Wild | 9.8% | 1.19 | 0.22 |
| | Domesticated | 10.9% | 0.49 | 0.57 |
| Soybean | Wild | 76% | 34.93 | 0.001 |
| | Domesticated | 75% | 30.39 | 0.001 |
| | Wild | 84% | 49.9 | 0.003 |
| | Domesticated | 74% | 198.05 | 0.001 |

### List of references of the studies analyzed in this work


1. Abera S, Shimels M, Tessema T, Raaijmakers JM, Dini-Andreote F. 2022. Back to the roots: defining the core microbiome of Sorghum bicolor in agricultural field soils from the centre of origin. *FEMS microbiology ecology* 98: fiac136.
2. Alegria Terrazas R, Robertson-Albertyn S, Corral AM, Escudero-Martinez C, Kapadia R, Balbirnie-Cumming K, Morris J, Hedley PE, Barret M, Torres-Cortes G, *et al.* 2022. Defining composition and function of the rhizosphere Microbiota of barley genotypes exposed to growth-limiting nitrogen supplies. *mSystems* 7: e0093422.
3. Chang C, Chen W, Luo S, Ma L, Li X, Tian C. 2019. Rhizosphere microbiota assemblage associated with wild and cultivated soybeans grown in three types of soil suspensions. *Archiv fur Acker- und Pflanzenbau und Bodenkunde* 65: 74–87.
4. Chang J, Costa OYA, Sun Y, Wang J, Tian L, Shi S, Wang E, Ji L, Wang C, Pang Y, *et al.* 2025. Domesticated rice alters the rhizosphere microbiome, reducing nitrogen fixation and increasing nitrous oxide emissions. *Nature communications* 16: 2038.
5. Coleman-Derr D, Desgarennes D, Fonseca-Garcia C, Gross S, Clingenpeel S, Woyke T, North G, Visel A, Partida-Martinez LP, Tringe SG. 2016. Plant compartment and biogeography affect microbiome composition in cultivated and native Agave species. *The new phytologist* 209: 798–811.
6. Cordovez V, Rotoni C, Dini-Andreote F, Oyserman B, Carrión VJ, Raaijmakers JM. 2021. Successive plant growth amplifies genotype-specific assembly of the tomato rhizosphere microbiome. *The Science of the total environment* 772: 144825.
7. Favela A, Bohn M, Kent A. 2022. N-cycling microbiome recruitment differences between modern and wild *Zea mays*. *Phytobiomes journal* 6: 151–160.
8. French E, Tran T, Iyer-Pascuzzi AS. 2020. Tomato genotype modulates selection and responses to root Microbiota. *Phytobiomes journal* 4: 314–326.
9. He X, Wang D, Jiang Y, Li M, Delgado-Baquerizo M, McLaughlin C, Marcon C, Guo L, Baer M, Moya YAT, *et al.* 2024. Heritable microbiome variation is correlated with source environment in locally adapted maize varieties. *Nature plants* 10: 598–617.
10. Kinnunen-Grubb M, Sapkota R, Vignola M, Nunes IM, Nicolaisen M. 2020. Breeding selection imposed a differential selective pressure on the wheat root-associated microbiome. *FEMS microbiology ecology* 96.
11. Leff JW, Lynch RC, Kane NC, Fierer N. 2017. Plant domestication and the assembly of bacterial and fungal communities associated with strains of the common sunflower, *Helianthus annuus*. *The new phytologist* 214: 412–423.
12. Liu F, Hewezi T, Lebeis SL, Pantalone V, Grewal PS, Staton ME. 2019. Soil indigenous microbiome and plant genotypes cooperatively modify soybean rhizosphere microbiome assembly. *BMC microbiology* 19: 201.
13. Miao M, Lankau R. 2023. Plant host domestication and soil nutrient availability determine positive plant microbial response across the Solanum genus. *Journal of experimental botany* 74: 1579–1593.
14. Oyserman BO, Flores SS, Griffioen T, Pan X, van der Wijk E, Pronk L, Lokhorst W, Nurfikari A, Paulson JN, Movassagh M, *et al.* 2022. Disentangling the genetic basis of rhizosphere microbiome assembly in tomato. *Nature communications* 13: 3228.
15. Pantigoso HA, Manter DK, Vivanco JM. 2020. Differential effects of phosphorus fertilization on plant uptake and rhizosphere microbiome of cultivated and non-cultivated potatoes. *Microbial ecology* 80: 169–180.
16. Pérez-Jaramillo JE, Carrión VJ, Bosse M, Ferrão LFV, de Hollander M, Garcia AAF, Ramírez CA, Mendes R, Raaijmakers JM. 2017. Linking rhizosphere microbiome composition of wild and

domesticated Phaseolus vulgaris to genotypic and root phenotypic traits. *The ISME journal* 11: 2244–2257.
17. Pérez-Jaramillo JE, de Hollander M, Ramírez CA, Mendes R, Raaijmakers JM, Carrión VJ. 2019. Deciphering rhizosphere microbiome assembly of wild and modern common bean (Phaseolus vulgaris) in native and agricultural soils from Colombia. *Microbiome* 7: 114.
18. Shenton M, Iwamoto C, Kurata N, Ikeo K. 2016. Effect of wild and cultivated rice genotypes on rhizosphere bacterial community composition. *Rice (New York, N.Y.)* 9: 42.
19. da Silva JL, Mendes LW, Rocha SMB, Antunes JEL, Oliveira LM de S, Melo VMM, Oliveira FAS, Pereira AP de A, Costa G do N, da Silva VB, *et al.* 2023. Domestication of Lima bean (Phaseolus lunatus) changes the microbial communities in the rhizosphere. *Microbial ecology* 85: 1423–1433.
20. Smulders L, Benítez E, Moreno B, López-García Á, Pozo MJ, Ferrero V, de la Peña E, Alcalá Herrera R. 2021. Tomato domestication affects potential functional molecular pathways of root-associated soil bacteria. *Plants* 10: 1942.
21. Tian L, Shi S, Sun Y, Tran L-SP, Tian C. 2020. The compositions of rhizosphere microbiomes of wild and cultivated soybeans changed following the hybridization of their F1 and F2 generations. *European journal of soil biology* 101: 103249.
22. Tronson E, Kaplan I, Enders L. 2022. Characterizing rhizosphere microbial communities associated with tolerance to aboveground herbivory in wild and domesticated tomatoes. *Frontiers in microbiology* 13: 981987.
23. Wipf HML, Coleman-Derr D. 2021. Evaluating domestication and ploidy effects on the assembly of the wheat bacterial microbiome. *PloS one* 16: e0248030.
